# HPCI: A Perl module for writing cluster-portable bioinformatics pipelines

**DOI:** 10.1101/408666

**Authors:** John M Macdonald, Christopher M Lalansingh, Christopher I Cooper, Anqi Yang, Felix Lam, Paul C Boutros

## Abstract

**Background:** Most biocomputing pipelines are run on clusters of computers. Each type of cluster has its own API (application programming interface). That API defines how a program that is to run on the cluster must request the submission, content and monitoring of jobs to be run on the cluster. Sometimes, it is desirable to run the same pipeline on different types of cluster. This can happen in situations including when:

- different labs are collaborating, but they do not use the same type of cluster
- a pipeline is released to other labs as open source or commercial software
- a lab has access to multiple types of cluster, and wants to choose between them for scaling, cost or other purposes
- a lab is migrating their infrastructure from one cluster type to another
- during testing or travelling, it is often desired to run on a single computer

However, since each type of cluster has its own API, code that runs jobs on one type of cluster needs to be re-written if it is desired to run that application on a different type of cluster. To resolve this problem, we created a software module to generalize the submission of pipelines across computing environments, including local compute, clouds and clusters.

**Results:** HPCI (High Performance Computing Interface) is a Perl module that provides the interface to a standardized generic cluster.

When the HPCI module is used, it accepts a parameter to specify the **cluster** type. The HPCI module uses this to load a driver HPCD∷<**cluster**>. This is used to translate the abstract HPCI interface to the specific software interface.

Simply by changing the **cluster** parameter, the same pipeline can be run on a different type of cluster with no other changes.

**Conclusion:** The HPCI module assists in writing Perl programs that can be run in different lab environments, with different site configuration requirements and different types of hardware clusters. Rather than having to re-write portions of the program, it is only necessary to change a configuration file.

Using HPCI, an application can manage collections of jobs to be runs, specify ordering dependencies, detect success or failure of jobs run and allow automatic retry of failed jobs (allowing for the possibility of a changed configuration such as when the original attempt specified an inadequate memory allotment).

## Background

Biocomputing pipeline programs are often written to be run on a computing cluster of some sort. We use the term **cluster** to refer not only to private collections of computers managed by control software for distributing jobs to those computers, but also to commercial cloud systems and their own control software. We also consider a single computer to be a degenerate form of cluster. As the amount of data and analysis involved continues to increase, so does the need for cluster-enabled pipelines.

Each type of cluster has its own application program interface (API) that is used to interact with the cluster in several ways. Most common of these is submitting a new job – including specification of resource requirements (*e.g*. memory, CPU, *etc.),* actions to be taken at job termination and dependencies amongst this job and others. Cluster APIs also support determination of the execution status of a submitted job, including distinguishing possibilities like the job is still running or that it has completed – normal termination with either an error status or successfully, or aborted by the cluster management software.

This API is often provided as a collection of utility programs that can be run manually or with a script. Often there is also a code library that can be used directly by programs as well.

It is often desirable to run the same program on multiple types of cluster. For example, when:

- different labs are collaborating, but they do not use the same type of cluster
- a pipeline is released to other labs as open source or commercial software
- a lab has access to multiple types of cluster, and wants to choose between them for scaling, cost or other purposes
- a lab is migrating their infrastructure from one cluster type to another
- during testing or travelling, it is often desired to run on a single computer instead of a full cluster

Unfortunately, each type of cluster has its own API - usually quite different from other cluster types. So, a program written to interact with one type of cluster must be significantly rewritten to interact with a different type of cluster. Thus, for a program to be able to run on multiple cluster type, you either have to recompile with different parameters, or change it in some other way to adapt to the type of cluster being used.

Regardless of whether the different labs are using the same type of cluster, different lab environments will have differences in the cluster node layouts to be used by the jobs and the “boilerplate” code that all software uses for the specific environment. For example, their nodes can be running different operating system variants requiring the programs use appropriate search paths for auxiliary programs; they can provide access to a choice of many different versions of programs and libraries but using differing mechanisms for the selection process.

## Implementation

### Solution

To deal with these issues, a Perl module HPCI [1] (High Performance Computing Interface) has been written that allows a program to be coded to run on a generic type of cluster. Additional modules are provided in the HPCD (High Performance Computing Driver) module hierarchy, that allow this generic cluster interface program to be executed on a specific cluster type by mapping the generic HPCI interface into the specific requirements of the selected cluster.

The HPCD driver module is loaded by HPCI automatically as it sets up its processing, based upon a parameter that specifies the cluster type to be used. A program using HPCI can be written to provide this parameter from the value given in a command line parameter or a configuration file. Thus, the same program can be re-run to use a different type of cluster just by providing a different value for the parameter and no change to the actual code.

Additionally, as HPCI starts up, it will automatically load a module named HPCI∷LocalConfig if it exists. Such a module can provide default settings that customize the HPCI activities to fit the local environment requirements.

The HPCI module is available as open source, and has been distributed on CPAN (the Comprehensive Perl Archive Network) which is used as the standard distribution mechanism for the vast majority of open source Perl modules. Documentation for HPCI can be displayed with the command “perldoc HPCI”.

### Generic HPCI Interface

The HPCI generic interface provides a program the ability to create a group that contains a number of stages.

A **stage** describes a single sub-program to be executed on the cluster.

The **group** manages the **stages**. It must be told of any execution order dependencies between stages. Attributes of the group specify the name of the group, control directory layout and logging for the group’s execution. The one required attribute is **cluster** - the type of cluster that will be used.

A group object has methods to create stages, to specify ordering dependencies between the stages and to execute the prepared group returning the completion status of the stages that were executed.

During execution, HPCI provides facilities to testing the success of the execution of each stage (with the ability to retry stages on failure), limit the number of stages being executed (both for cluster limits and to meet dependency ordering requirements) and to determine whether individual stage failures necessitate aborting the remainder of the group execution.

An HPCI internal object **run** is used for each attempted execution of a **stage**.

The **stage** method used to create a stage can be provided with a number of attributes. These specify:

- the name for the stage (required, must be unique)
- the command to be executed
- the resources that the stage requires (memory, execution time, cpu cores, *etc*.)

These resources can be specified as soft or hard. The difference between soft and hard lies in the action taken when the resource limit is exceeded. A soft limit causes the program to be sent a signal, giving it a chance to shut down cleanly; a hard limit exception causes the program to be terminated. The driver for any particular type of cluster may support a subset of these capabilities and should map the generic request into the closest fit available from that actual cluster type.

There are different actions that can be taken on completion of a stage. A program can provide call-back functions to determine whether the stage completed successfully - although normally, the HPCI method of checking the exit status of the stage is sufficient. If a stage fails to execute properly, there are facilities for deciding whether to retry the stage - either using different memory resource requirements if the stage was killed for running out of memory, or running the stage the same way when there is the possibility that some compute nodes on the cluster could fail when others would succeed. If a failed stage is not being retried, there are options to decide whether to stop executing any other stages - the default is to continue to execute stages that are not dependent upon the failed one.

### Configuration Considerations

There are a number of different ways that a program may need to be controlled to deal with moving from one run environment to another.

- Different types of underlying cluster software will require mechanisms for mapping the generic HPCI definitions to those used by the job dispatching functions of the cluster; there are often similarities between these mapping mechanisms that make it useful for them to use a common method with a few parameters specific to the cluster type.
- Different labs will have their own local requirements for such issues as setting the environment for cluster jobs, choosing (as part of the run configuration) the version of libraries and programs to be used.
- Individual programs may wish to provide common attribute settings to all or most of the stages that they run.
- Individual programs may wish to provide some driver-specific attributes to be used depending upon the actual cluster that is selected for an individual run.

The objects used by HPCI (group, stage and internal objects) are implemented using roles. Some of the roles are always used, but some can be specified by the driver or by a local configuration module.

### Configurability to Cluster Type

Drivers can list additional roles to be used explicitly, as a driver is itself a collection of roles that are merged with the generic object types.

However, a driver generally provides or overrides methods in the objects to carry out the driver-specific functions.

For example, there is a generic mechanism that will map driver-specific resource names to the generic HPCI resource names, allowing a driver to accept the names commonly used by its own cluster type. This means existing programs can be more quickly converted from the native cluster coding style to HPCI (albeit with reduced flexibility to run on other cluster types until those names are changed to the generic ones).

As another example, the generic HPCI code assigns default values for resources and retry_resources, but calls a method to get the values to be assigned. The generic value method provides an empty list of values, but a driver can replace that method with another which provides a different list.

### Configurability to Local Environment

When the HPCI module is loaded, it tries to load a module named **HPCI∷LocalConfig**. If such a module is not found, there are no ill effects. No such module is provided in the HPCI distribution. An organization, however, can write such a module to provide any customization required for the local environment. This module can use the following HPCI class methods to install these customizations.

- **add_extra_role** This class method will accept the names of roles to be included into group or stage objects. They can be made conditional on the cluster type, or provided for all cluster types. The parameter is a hash with keys that are either a cluster type or “ALL”. The value is a hash with a keys of either **stage** and/or **group**. The value for these keys is a list of additional roles to be merged into the specified object type. The values for “ALL” are merged in first, then the values for the actual cluster type being used are merged in thus taking precedence if they overlap.
- **add_default_attrs** This class method will accept a hash containing default attribute parameters to be provided to the creation of a group.

Some sub-roles can also be configured. For example, the **HPCI∷ScriptSource** module provides the ability to generate a script to be executed when a stage is run. It provides an array:

- **HPCI∷ScriptSource∷source_commands** This array can be loaded with text lines to be pre-pended to the script before the command that is the body of the stage.

### Available Drivers

The HPCI package includes a single driver, for a cluster type called **uni**. The **uni** driver does not actually use a cluster of multiple machines. Instead it runs stages by using the Unix **fork** mechanism and runs the stages on the same machine as the parent program. This driver is used by default for running the HPCI test suite on installation. That ensures that HPCI can be installed even if there are no HPCD driver packages yet installed. The **uni** driver can also be useful for programs that use HPCI - both for programs that are using small enough data sets that they do not need to be run on an actual cluster and as a fall-back to run a program even if the cluster is not available for some reason.

HPCI [1] is available from **CPAN**.

In addition to the **uni** driver that comes with HPCI, there are drivers separately available for two types of cluster.

**HPCD∷SGE** [2] provides the driver for SGE (Sun Grid Engine) clusters.

**HPCD∷SLURM** [3] provides the driver for Slurm clusters (SLURM was originally an acronym for Simple Linux Utility for Resource Management, but the acronym is only referred to as having historical meaning).

These two drivers were chosen because of the needs of our lab (SGE) and our first collaborator to require a different cluster (Slurm). Additionally, they both are widely used, in particular, “Slurm is the workload manager on about 60% of the TOP500 supercomputers, including Tianhe-2 that, until 2016, was the world’s fastest computer.” [4]

In the future, there will be additional drivers from the authors. Because **HPCI** is open source, people from other labs can write drivers for their own cluster types without relying on the authors to write a driver for them. The authors encourage such independent driver development and are happy to assist in such efforts.

## An Example

Let us show a very simple pipeline. This pipeline takes two files, *XY.normal.vcf* and *XY.tumour.vcf,* runs each of them through a filter program *filter,* generating the output files *XY.normal.filter.vcf* and *XY.tumour.filter.vcf.* Those steps can be done simultaneously. Then an analysis program *analyze* is used to determine the differences between the normal and tumour data, generating a report file *XY.rpt* and a graph file *XY.pdf.* (The name of the study **XY** is provided to the program as a parameter in this example, although in practice it would more often be provided along with many other parameters in a configuration file.)

A program to do this directly using an SGE cluster (omitting a lot of error detection logic and some necessary subroutine functions) would look like:

~~~
my $study_name = shift;
my %running_processes;
for my $type (qw[normal tumour]) {
    my $fileprefix = “$study_name.$type";
    my $thispid = forkcmd(“qsub -N $fileprefix “
       . “-sync y -cwd”
       . “filter -i $fileprefix.vcf”
       . “-o $fileprefix.filter.vcf”
       );
   $running_processes{$thispid} = 1;
   }
while(keys %running_processes) {
   my $pid = waitpid −1,0;
delete $running_processes{$pid} if exists $running_processes{$pid};
   }
my $analysis_process = forkcmd(“qsub “
  . “-N $study_name.analysis “
  . “-sync y -cwd “
  . “analyse “
      . “-n $study_name.normal.filter.vcf ”
      . “-t $study_name.tumour.filter.vcf ”
      . “-r $study_name.rpt ”
      . “-p $study_name.pdf”
  );
1 until $analysis_process eq waitpid −1,0;
# functions not shown here:
# waitpid should be wrapped with a check function that
# - calls the waitpid built-in as above
# - abort if there are no child processes running
# forkcmd
# - fork a child process to execute a provided command
# - return the child’s process id
~~~

This code would only work on an SGE cluster, of course.

Using HPCI the code would be:

~~~
use HPCI;
my $study_name = shift;
my $cluster = shift // ‘SGE’;
my $group = HPCI->group(cluster => $cluster, name => $study_name);
my @filter_stages;
for my $type (qw[normal tumour]) {
    my $fileprefix = “$study_name.$type";
    my $this_stage_name = “$study_name.$type.filter";
    $group->stage(name => “$this_stage_name",
        command => “filter -i $fileprefix.vcf -o $fileprefix.filter.vcf”
       );
    push @filter_stages, $this_stage_name;
    }
my $analysis_stage_name = “$study_name.analyse";
$group->stage(name => $analysis_stage_name,
    command => “analyse “
       . “-n $study_name.normal.filter.vcf “
       . “-t $study_name.tumour.filter.vcf “
       . “-r $study_name.rpt “
       . “-p $study_name.pdf”
   );
$group->add_deps(pre_reqs => \@filter_stages, dep => $analysis_stage_name);
$group->execute;
~~~

The code using HPCI is shorter. It does not have to manually deal with coordinating dependencies between stages that need to be run in constrained order. It has all of the error checking that the original code does not (provided automatically by HPCI). While an SGE-specific module package could certainly provide the same ease and error checking, the HPCI code is not tied to the SGE cluster - it could run on a different type of cluster simply by providing (in this example’s case) a second command line argument.

## Related Work

There are library modules for many languages that allow easy definition of parallel programming, but most of these are aimed at using multiple processes on the same processor (with the separate jobs spawned off using fork on modern Unix processors and corresponding system interfaces of other operating systems). Such modules date back many decades.

However, for parallel processing on collections of computers most library modules only provide an interface to a specific type of cluster.

There is a C library, DRMAA [5], that provides a common interface to a number of cluster types. HPCI uses a Perl module that interacts with DRMAA for the SGE cluster and may do so for other cluster types in the future (the Slurm cluster has a DRMAA interface but the HPCD∷SLURM driver is not using it at this time. (The Perl Schedule∷DRMAA package that interfaces to a DRMAA binary library has only ever been used with the SGE DRMAA interface; we have not gotten it to link correctly with DRMAA that comes with Slurm.)

There is a Python module Ruffus [6] which is capable of running jobs on clusters that provide a DRMAA library interface. Since our lab uses Perl for virtually all of our control scripts, this was not considered a good alternative for our purposes (in fact, we only noticed it while doing a survey of related work for this article).

## Discussion

The design of HPCI was inspired by the extremely widely used DBI/DBD [7] modules (Data Base Interface / Data Base Driver) that are commonly used for database programming in Perl. The DBI module provides an interface to databases, and the DBD modules get selected automatically to connect DBI-based programs to actual databases (such as Oracle, DB2, Sybase, *etc.*).

Like DBI, HPCI does not force users to write their code portably. HPCD drivers accept driver-specific names for resources, and they can have limitations forced upon them by the underlying cluster API not perfectly matching the HPCI generic interface. For example, the SGE cluster supports hard and soft limits on resources, but a stage may not specify both. That is conceptually useful (give the stage program a warning signal, allowing it to clean up and terminate in a controlled fashion, while still forcing the stage to terminate if it continues to further exceed the limit). The SGE API simply applies the last limit specified and ignores the previous ones. So, the HPCD∷SGE driver provides only the soft limit and drops the hard limit (thus the stage program can either catch the signal and clean up correctly, or not catch the signal at all and be killed immediately by it).

So, it still requires a degree of careful coding by the user programmer to cast their pipeline in a way that is truly portable to multiple cluster types.

HPCI provides mechanisms for specifying file requirements for stages - when a cluster type that does not have a shared file system is encountered, using these mechanisms will become critical.

However, at the current time, all of the usage of HPCI has been done on clusters that provide a shared file system accessible to all of the nodes, so programs have being written without specifying file requirements and so they will need updates targets missing such a shared file system.

## Results

The HPCI and some HPCD modules are available as open source to be used by anyone, including all research institutions.

## Conclusion

HPCI is a valuable module for allowing programs to be written with portability across multiple types of cluster system, without losing the ability to take full advantage of specific cluster systems when needed. This allows collaboration and validation to take place in labs which use different cluster mechanisms, as well as allowing the software component results of research to be released as open source in a form that can be used by many research institutions without the need for major re-coding.

## Declarations

## List of Abbreviations

API: Application Programming Interface
CPAN: Comprehensive Perl Archive Network
DBD: Data Base Driver
DBI: Data Base Interface
HPCD: High Performance Computing Driver
HPCI: High Performance Computing Interface
SGE: Sun Grid Engine
SLURM: Simple Linux Utility for Resource Management

## Ethics Approval and consent to participate

Not applicable

## Consent for publication

Not applicable

## Availability of data and material

Data sharing not applicable to this article as no datasets were generated or analysed during the curtrent study.

## Author’s contributions

This paper was initially drafted by John Macdonald. It was edited and approved by all of the authors. Dr. Paul Boutros supervised the research. The code for the main module (HPCI) and the two initial drivers (HPCD∷uni and HPCD∷SGE) were written by John Macdonald, by rewriting modules written for SGE-specific cluster usage (written by Christopher Cooper, Christopher Lalansingh, and others) to separate cluster-specific and cluster-generic components. The code for the HPCD∷SLURM driver was written by Anqi Yang based upon the HPCD∷SGE driver. Test modules were written by John Macdonald and Felix Lam.

## Acknowledgements

Special thanks to the members of the Boutros Lab team for extensive use and feedback for the HPCI and HPCD∷SGE modules in a wide range of pipelines.

